# DrugProtAI: A guide to the future research of investigational target proteins

**DOI:** 10.1101/2024.11.05.622045

**Authors:** Ankit Halder, Sabyasachi Samantaray, Sahil Barbade, Aditya Gupta, Sanjeeva Srivastava

## Abstract

Drug design and development are central to clinical research, yet ninety percent of drugs fail to reach the clinic, often due to inappropriate selection of drug targets. Conventional methods for target identification lack precision and sensitivity. While various computational tools have been developed to predict the druggability of proteins, they often focus on limited subsets of the human proteome or rely solely on amino acid properties. To address the challenge of class imbalance between proteins with and without approved drugs, we propose a novel Partitioning Method. We evaluated the druggability potential of 20,273 reviewed human proteins, of which 2,636 have approved drugs. Our comprehensive analysis of 183 features, encompassing biophysical and sequence-derived properties, achieved a median AUC of 0.86 in target predictions. We utilize SHAP (Shapley Additive Explanations) scores to identify key predictors and interpret their contribution to druggability. We have reviewed and evaluated 688 investigational proteins from DrugBank (https://go.drugbank.com/) using our tool, DrugProtAI (https://drugprotai.pythonanywhere.com/). Our tool offers druggability predictions and access to 2M+ publications on drug targets and their effects, aiding in the selection of target proteins for drug development. We believe that insights into key predictors will significantly advance drug development and propel the field forward.

## Introduction

Drug design and development have been arduous tasks, yet a staggering 90% of drugs fail to reach the clinic^1^. The ability to predict whether a protein can effectively bind to small molecules or drugs has long been a key focus of interest for various structural and computational biologists^2^. The sole dependence on the 3D structure of proteins and the limited structure availability always hindered the process of accurate target identification^3^. The advent of deep learning models integrated into accurate identification of the 3D structure of proteins^4,5^ has bolstered the field to a great extent. Nevertheless, druggability prediction remains limited due to the failure to account for all associated physicochemical and biological properties. A substantial proportion of drugs fail to accomplish the goal of implementation in medical practice due to their targets being considered undruggable^6^. It is also essential to comprehend the distinction between druggability and ligandability. Druggability pertains to the therapeutic action of a drug whereas ligandability refers to the ability to bind to small molecules or ligands^7^. The assessment of the drug likeness properties of the proteins^8,9^ and the association of the human genes encoding druggable proteins^7,10^ have remained subjects of continued scientific investigation.

The last decade has seen an unprecedented increase in studies that have employed machine learning (ML)-based computational approaches to identify druggable proteins, with the sole aim of complementing traditional and experimental methods. Shoombuatong, *et. al* surveyed and highlighted different tools, algorithms and deployed webservers spanning this space, yet the full realization of maximal accuracy in uncovering the druggability potential of protein targets remains elusive^11^. Major concerns of the developed tools and algorithms towards the identification of druggable proteins have been raised and rightly pointed out by Charoenkwan *et. al* which led to the development of a new tool, SPIDER^12^ (http://pmlabstack.pythonanywhere.com/SPIDER). However SPIDER also suffers from certain limitations, such as the training set comprises only 2,543 proteins. Additionally, the model is trained solely on features derived from amino acid sequences and their properties. The approach also lacks reference to key biophysical predictors that are fundamental in predicting protein druggability. Another recent tool, DrugnomeAI (https://astrazeneca-cgr-publications.github.io/DrugnomeAI/), which is based on a stochastic semi-supervised machine learning framework predicts the druggability of protein-coding genes by integrating gene-level data resources, but it lacks emphasis on features specific to proteins^7^. Furthermore, these tools fail to provide comprehensive information related to recent literature regarding proteins as candidate drug targets for various diseases.

To overcome the aforementioned drawbacks, we developed an advanced computational solution to address the significant class imbalance, resulting in the creation of a tool called the DrugProtAI. DrugProtAI has been designed by modelling almost the entire set of human proteome, thereby circumventing the limitations associated with focusing on only a subset of human proteins, as seen in other tools and algorithms. Secondly, the features have been extracted from UniProt^13^ and the model is based on ten major types of features, including physicochemical properties, grouped dipeptide composition (GDPC) encodings, properties derived from protein-protein interactions(PPIs), counts of post-translational modification residues (PTM counts), subcellular localizations, domains, and sequence-encoded latent values (**Supplementary Table 1**). This approach effectively mitigates the constraints inherent in relying solely on amino acid properties, as observed in numerous other tools. Thirdly, the partitioning method has been employed to create an equal representation of druggable and non-druggable proteins for the training of the model, which was subsequently evaluated using an independent dataset. The models have been evaluated using various random seed selections to ensure an unbiased estimation of the reported final metrics. Finally, gradient boosting and random forest methods have been integrated with the partitioning approach to predict the druggability of the proteins. Moreover, SHAP scores have been utilized to delineate the principal predictors and provide interpretable insights, thereby aiding subsequent researchers in the evaluation of the protein prior to its selection as a drug target. We propose two different modellings for druggability prediction – namely Partition Ensemble Classifier(PEC) and Partition Leave-One-Out Ensemble Classifier (PLOEC). PLOEC ensures that the weight of the partition in which the protein is used as a train set is nullified to retrieve accurate results during druggability evaluation.

The data from Drug Bank (https://go.drugbank.com/), as of March 16, 2024, included 688 investigational target proteins associated with various drugs in the context of different diseases. All the investigational proteins have been tested using DrugProtAI to assess their potential to be druggable. Out of these proteins, 10 have received approval for a drug as of August 10, 2024, and an additional 13 proteins have also received approval for new drugs **(Supplementary Table 2)**. All of these were evaluated using the method to assess the tool in the form of blinded validation. These proteins were also tested using existing benchmarking tools and compared with DrugProtAI. Our modelling strategy provides more robust results compared to others (**Table 1)**.

**Table 1.** | Comparison of our Partition Models (PEC and PLOEC) against SPIDER. Predictive accuracy on the newly approved druggable proteins using the amino acid sequence based SPIDER method and our PEC and PLOEC ensemble classifiers using XGBoost and Random Forest.

| Method | Proteins : Investigational to<br>Approved [10 proteins]<br>(Accuracy in %) | Proteins : Non-Druggable to<br>Approved [13 proteins]<br>(Accuracy in %) | Proteins : Investigational to<br>Approved [23 proteins]<br>(Accuracy in %) |
| --- | --- | --- | --- |
| DrugProtAI (XGB) | 80 | 84.62 | 82.61 |
| DrugProtAI (RF) | 70 | 76.92 | 73.91 |
| SPIDER<br>(Charoenkwan et al., 2022) | 70 | 53.85 | 60.87 |

Understanding the features that resemble closeness to proteins with druggable properties, along with further assessment using DrugProtAI, will help investigators make informed decisions regarding the selection of specific targets. The advent of high-throughput omics technologies can pave the way for the identification of a large number of drug targets^14^, and DrugProtAI can be used as a complementary tool in the drug discovery process. The search for recent literature on investigational targets is often cumbersome; therefore, DrugProtAI has integrated a feature for retrieving publications that provide supporting evidence for the protein as a drug target. We believe that the easy-to-use web interface of DrugProtAI, will be a small leap towards dispelling the uncertainty surrounding druggable proteins, potentially saving a significant amount of time and money in the future.

## Results

We intend to discover and interpret key predictors contributing to protein druggability and numerically capture the probability of a protein being druggable.

### Feature Extraction and Engineering

We extracted comprehensive protein information from the UniProt^13^ database and cross-referenced it with DrugBank, resulting in a broad classification of the proteome into three categories simply based on their association with drugs – Druggable, Investigational and Non-Druggable. With approximately 13.1% in the druggable and 83.6% in the non-druggable classes (**Fig. 1a**), the significant skewness poses critical challenges in analysis. From the extracted protein data, we derived 183 features relevant to the druggability potential of a protein and categorized them into ten distinct feature classes. The distribution of features within each category is illustrated using a radar plot **(Fig. 1b)**. A complete list of features within each group is provided in **Supplementary Table 1**. The following is a detailed description of each feature class.

**Fig. 1.**
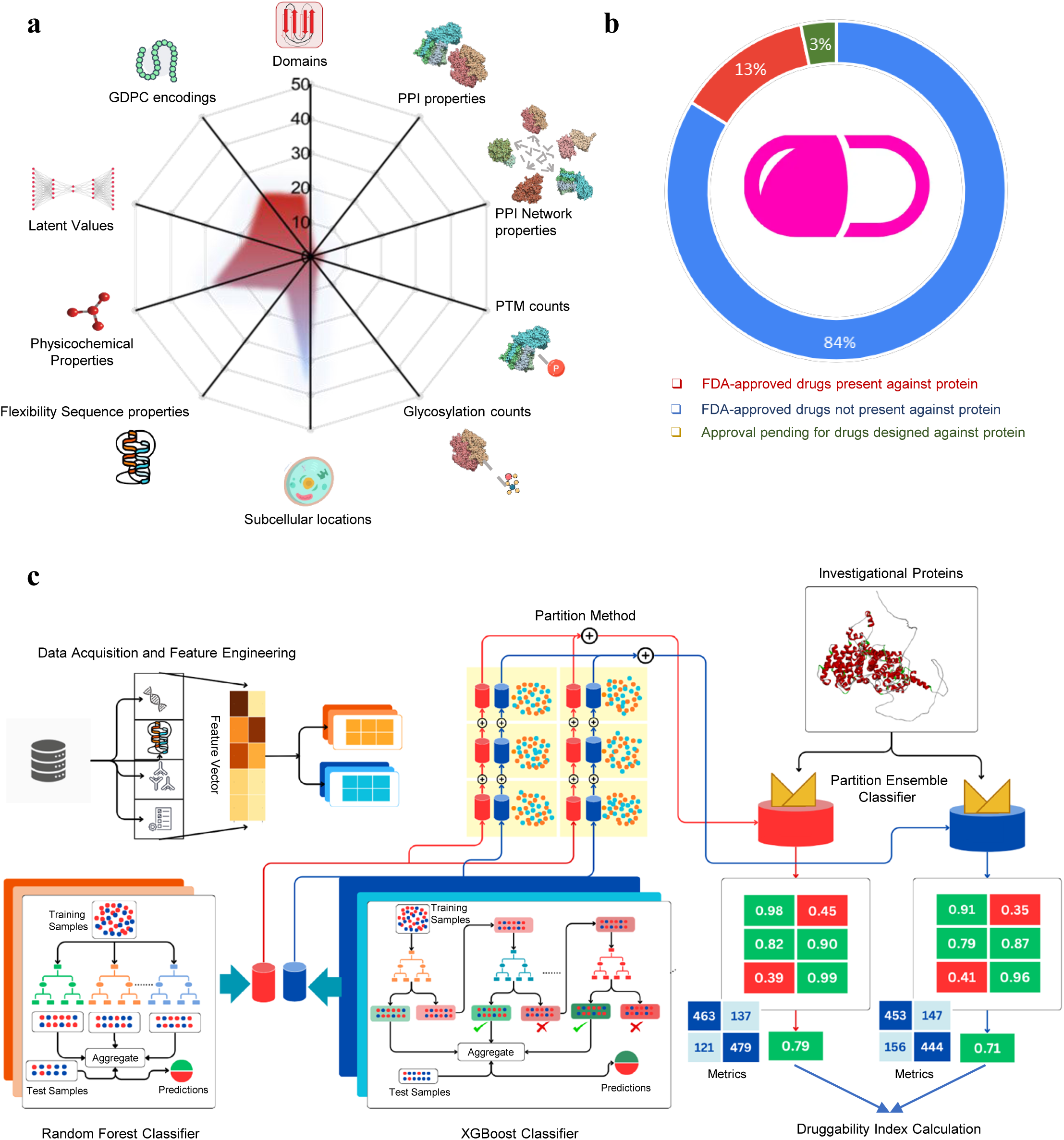
| Illustration of the Extracted Dataset and Druggability Prediction Model Development Workflow. **a**, We utilized a comprehensive feature set that includes PPI properties (15%), PTM counts (10%), glycosylation counts (8%), subcellular locati ons (12%), flexibility sequence properties (10%), and other domain-specific features, forming a robust input for model training. **b,** The dataset, consisting of 20,343 proteins, is categorized into 83.47% non-druggable, 13.13% with approved drugs, and 3.39% drugs pending approval. **c,** The non-druggable proteins are further divided into six partitions, which are then combined with the druggable samples to create balanced partitions with equal distribution, ensuring a non-skewed dataset . The model development workflow involves training ensemble classifiers, including Random Forest and XGBoost, on these balanced subsets. Each model undergoes validation and testing on an independent protein sample set, with a majority voting mechanism employed to integrate results, ensuring robust and accurate predictions. The final druggability index and metrics are calculated by averaging predictive scores from out-of-bag sets across the ensemble models, leading to reliable classification outcomes

Domain (20 features): Domain information was extracted from the UniProt^13^ features JSON. We selected the 20 most frequent domains across the entire dataset. These features capture the number of occurrences of the top 20 domains in the protein sequence as evidenced from published studies. Protein domains are indicative of its function, structural integrity and presence of potentially amenable sites for drug binding.

Protein-Protein Interaction (PPI) Properties (4 features): PPI information was extracted from the Binary Interactions section of UniProt^13^. These properties capture the number of binary interactions, xenobiotic interactions and the number of supporting published studies. PPIs reveal the role of a protein in biological communication pathways.

PPI Network Properties (7 features): Using the extracted PPI properties, we constructed a network represented as a graph with undirected edges and corresponding edge labels. An edge between proteins *P_i_* and *P_j_* indicates their interaction, with an edge label denoting whether the interaction is xenobiotic or binary. Features in this category include network properties such as node degree, average degrees of neighboring proteins, and the sizes of strongly connected components. PPI Network properties reveal interconnected pathways for therapeutic strategies.

Post-Translational Modification (PTM) Counts (5 features): PTM counts were extracted from the UniProt^13^ Features JSON. These features include counts for glycosylation, cross-linking, modified residues, signal peptides, and disulfide bonds. PTM’s influence a protein’s druggability by affecting its stability, activity, and interaction capabilities.

Glycosylation Counts (6 features): Glycosylation counts were extracted from the UniProt^13^ Features JSON. These features include various types of glycosylation, such as O-linked, N-linked, S-linked, C-linked, O-alpha-linked, and N-beta-linked. Glycosylation can impact protein-ligand interaction and cellular localization.

Subcellular Locations (SCLs) (50 features): Information about SCLs were extracted from the Subcellular Locations section in UniProt^13^. These are grouped into 50 major categories, and then binary features are created to indicate the presence or absence of specific SCL within a protein. SCLs are indicative of a protein’s functional context and localization specific interaction.

Flexibility Sequence Properties (FSPs) (14 features): FSPs were calculated from amino acid sequences of proteins using Vihinen’s method^15^. These features describe the statistical properties of the protein sequence. FSPs are indicative of protein’s inherent flexibility and structural dynamics.

Physicochemical Properties (PCPs) (32 features): PCPs include amino acid composition, molar extinction coefficients, secondary structure content, sequence length, molecular weight, grand average of hydropathy (GRAVY), isoelectric point, instability index, charge at pH 7, and aromaticity, all derived from amino acid sequence of proteins using the BioPython package^16^ PCPs are indicative of a protein’s behavior in biological systems like solubility, protein folding, stability and hydrophobicity.

Latent Values (20 features): Latent Values were extracted from the latent intermediate layer (size = 20) of an autoencoder trained on protein sequences. These were designed to reconstruct the input sequence in a traditional encoder-decoder architecture, capturing the latent features crucial for entire protein representation.

Grouped dipeptide composition (GDPC) Encodings (25 features): GDPC encodings are 25 numerical values corresponding to dipeptide pairs (x,y) in sequence where x and y belong to residue categories such as aliphatic, aromatic, positively charged, negatively charged, or uncharged. The feature space for GDPC was formulated as provided in the study by Sikander et. al^17^

Each of these feature classes offered distinct insights into the protein properties potentially contributing to its druggability.

### Evaluating Partition Method

To assess the effectiveness of the proposed Partition based modeling (**Fig. 1c**), we partition the majority class (non-druggable train set) into eight partitions, each comprising approximately 2,044 proteins. Subsequently, we trained eight models on each of the non-druggable partition sets against the entire druggable train set, which comprises 2,036 proteins, therefore the class imbalance effects are mitigated. The overall accuracy and individual accuracies on the druggable and non-druggable test sets were included in the assessment measures. For both XGBoost (**Supplementary Table 3a**) and Random Forest (**Supplementary Table 3b**), we also included the performance metrics for each individual model in addition to the overall PEC model. To ensure variety in the selection of test data and training data partitions, the assessment was carried out using 20 distinct random seed configurations, simulating a cross-validation situation. The provided metrics are presented as mean values together with their variance across simulations. Interestingly, the PEC model’s overall accuracy was around two percentage points greater than the average accuracy found in all divisions. This result validates our hypothesis that the collective performance of the various models produces better results than the mean performance of the individual models. This confirms that our partitioning technique functions as intended. We also experiment with a Genetic Algorithm using Roulette Wheel Selection for feature selection on the XGBoost classifier in order to improve the computational efficiency and predictive power of our model (**Supplementary Fig. 1**). This allowed us to reduce the number of features from the original 183 to an average of roughly 85 across eight partitions, each of which contained 2036 proteins in the druggable train set. Additionally, we provided metrics in (**Supplementary Table 3c**) for the application of GA on XGBoost Classifier by the PEC technique. Its overall forecast accuracy was almost 77%, which was comparable to the performance of the other models. The PEC modeling has proven to be a reliable and robust strategy for classification of proteins based on druggability. We further examined the feature relevance using this PEC framework to get insights into the key predictors influencing druggable tendency of proteins, thereby improving feature selection and contributing to interpretability of the predictive model.

### Feature Improvement Metrics and Interpretability of Partitioning Models

To interpret key features responsible for contributing to druggability prediction of proteins, we calculated feature scores by averaging the individual importance scores from each of the models in our partition method (**Supplementary Table 4a** for XGBoost, **Supplementary Table 4b** for Random Forest). The features were then ranked based on their importance. Averaging ensures that characteristics consistently performing well across partitions are given the appropriate weight while also mitigating any biases introduced by specific partitions. This also prevents any single partition from having an excessively large impact on the final ranking. Additionally, SHAP values provided an interpretable understanding of each feature’s contribution to druggability classes. For each feature in every partition and for each sample, SHAP values were computed. These SHAP scores were then averaged across samples and partitions to gain insights into interpretability on both a local and global scale (**Fig. 2a**). The test accuracy and partition average feature scores improvements as the number of top features increasing from K = 1 to K = 183 is illustrated in **Fig. 2b**. The XGBoost model achieves maximal test accuracy of around 78% with 86 features (**Supplementary Table 5a**) whereas Random Forest achieves its maximum at a lower accuracy of around 76% at 125 features (**Supplementary Table 5b**). Because of the sensitive boosting method of XGBoost, which prioritizes distinct subsets of data in each partition, it has larger variance between partitions than Random Forest, as demonstrated by the Partition Average feature scores in decreasing order. However, greater stability across partitions is achieved by Random Forest’s ensemble technique, which averages data from several trees.

**Fig. 2.**
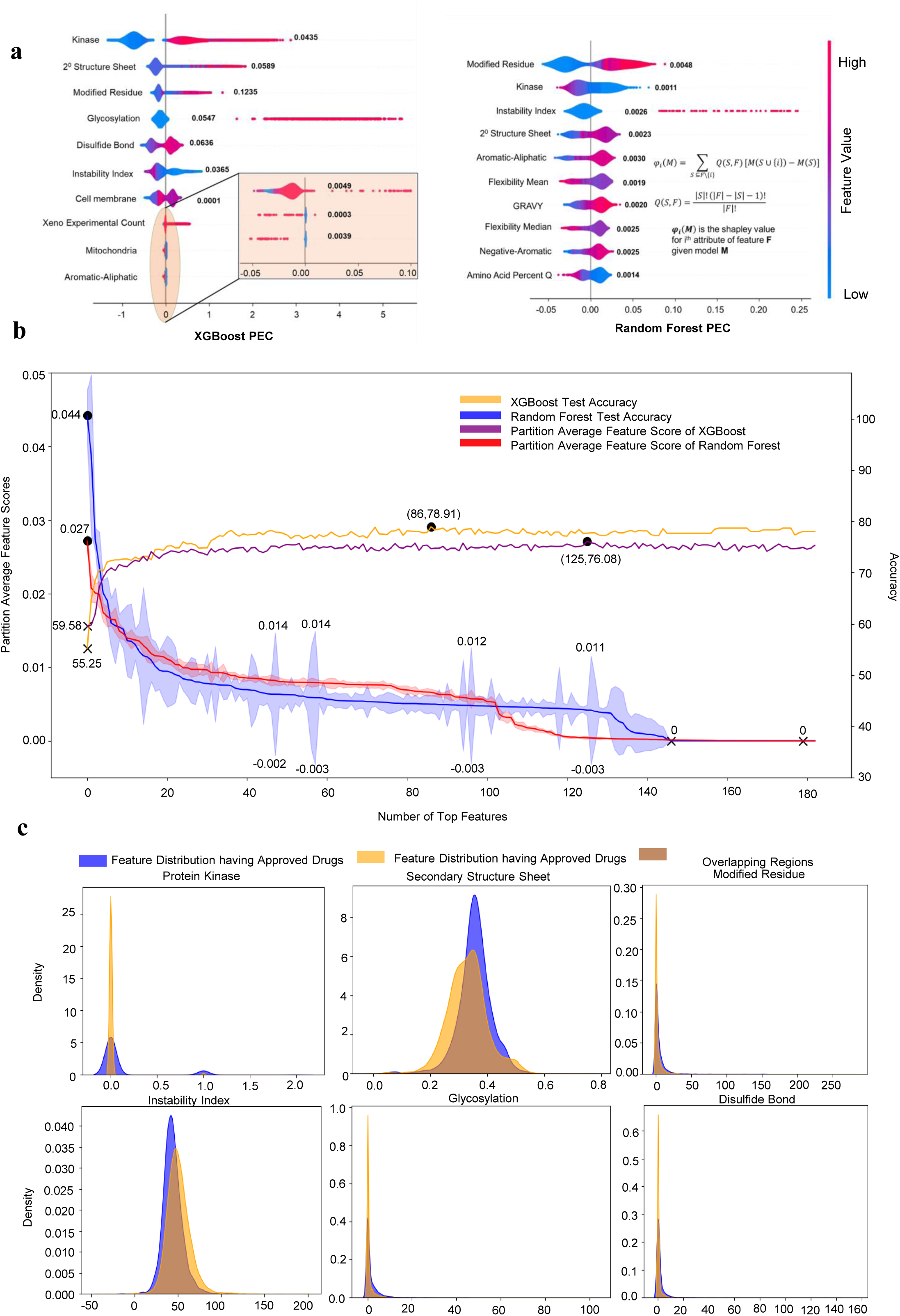
| Feature Scores, Feature Improvement Metrics and Distribution Plots of Top Features. **a**, Summary plots show SHAP values of the top 10 features selected based on Partition Average scores. The models used are XGBoost PEC and Random Forest PEC. The SHAP scores are calculated on a small representative subset of the training dataset itself. These plots contribute to the model’s interpretability of how feature values affect the druggability of the protein **b,** (Left Axis) Partition Average Feature Scores for both XGBoost and RF models in decreasing order. The curve also displays the variance in feature scores across individual partition models. (Right Axis) Improvement in test accuracy as more top features (from K=1 to K=183) are included. **c,** Demonstrates the data distribution plots for some of the top features from our Partition based ML modelling framework.

The top 10 features influencing druggability predictions are shown in **Fig. 2a**, where the SHAP summary plots for the Random Forest and XGBoost models are displayed. These plots share several important features, such as kinase, modified residue, instability index, and secondary structure sheet, underscoring their significance for both algorithms. Positive numbers push in the direction of the prediction being druggable, whereas negative values indicate the opposite. The x-axis represents the strength and direction of feature influence. The feature values are represented by color in the graphs. Blue denotes lower feature values, whereas Red implies higher feature values. Notably, proteins with greater counts of glycosylation or kinases make a significant and beneficial contribution to druggability. Higher feature values, on the other hand, have the reverse effect on druggability when it comes to flexibility factors like mean and median. In contrast, flexibility features such as mean and median behave opposite, with higher feature values contributing negatively to druggability. Features such as secondary structure sheets, modified residue counts, disulphide bond counts, presence of cell membrane, average hydrophobicity(GRAVY) contribute positively to druggability with higher feature values. There are several features where Random Forest and XGBoost do not show agreement in interpretation. The observations made by the XGBoost partitioning model align closely with findings from literature and distribution plots.

We sought to interpret which feature group contributes the most towards the prediction of protein druggability. For every feature group, the total contribution is calculated as the sum of feature importance scores of each constituting feature (**Supplementary Fig. 2**). Both models portray agreement in labeling Physicochemical Properties as the topmost contributing group. Features such as secondary structure sheet, instability index, grand average of hydropathy, and glutamine percentage in the amino acid sequence emerged as top features across both models. GDPC encodings and subcellular locations also played a major role in druggability prediction. The distribution pattern of some of the top features like secondary structure sheets, and instability index is also found to be quite distinct (**Fig. 2c** & **Supplementary Fig. 3**). The key predictors from our partition modeling can be considered reliable indicators due to their notable distinctions and thus serve as strong candidates for further investigation in drug repurposing strategies.

### Druggability Index and Blinded Validation

The model deployed in the tool available at https://drugprotai.pythonanywhere.com/, was trained on the entirety of 2636 druggable and 16949 non-druggable proteins, excluding investigational proteins from the training phase. To achieve this, the non-druggable set was divided into six partitions 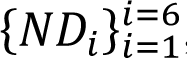, and 6 ensemble models 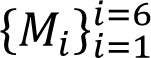 were trained, each using a partition of the non-druggable set (∼2824 proteins) against the entire druggable set (|D| = 2636 proteins).

We calculated the Druggability Index(DI) for the 688 investigational proteins, which were unseen during the training, using the PEC model. We also calculated DI for 16949 non-druggable proteins, using PLOEC model. These DI values are provided in **Supplementary Table 6a** for XGBoost PLOEC and **Supplementary Table 6b** for Random Forest PLOEC. DI can be considered as a measure of how strongly a protein exhibits druggable tendencies. It must be noted that over time, proteins may transition from one category to another, as new drugs are being investigated and approved. Our categorization was based on available drug information extracted from DrugBank as of 16th March 2024. All proteins which have moved from the investigational and non-druggable classes to higher categories, as of data extracted from 10th Aug 2024 are listed (**Supplementary Table 2)**. 3 proteins have entered the investigational category whereas 23 new proteins now have FDA approved drugs. This clearly reflects the pace in drug discovery research.

The XGBoost-based PEC/PLOEC models generated DI demonstrated higher accuracy in comparison against Random Forest. 19/23 druggable proteins were predicted correctly as druggable by our XGBoost based PEC and PLOEC model. This serves as a potential blind validation of our methods, with XGBoost achieving 82.6% accuracy and Random Forest achieving 73.9% accuracy (**Table 1**). Notably, Random Forest performs poorly compared to XGBoost, as was also observed in the behavior of the respective key predictors using SHAP scores. High druggability index scores indicate a greater likelihood of a protein’s potential to be targeted by a drug. Our scores can help identify potential targets for drug development based on their roles in disease progression. We observed 12 proteins in the investigational set that are predicted to have a DI > 0.99 by our XGBoost PEC model (**Supplementary Table 7a),** 2 of which have been already approved. Illustrative examples of how our method performs with investigational, druggable, and non-druggable proteins are presented (**Fig. 3a & Fig. 3b)**. We also provide DI scores of Random Forest PEC model on investigational proteins at **Supplementary Table 7b**.

**Fig. 3.**
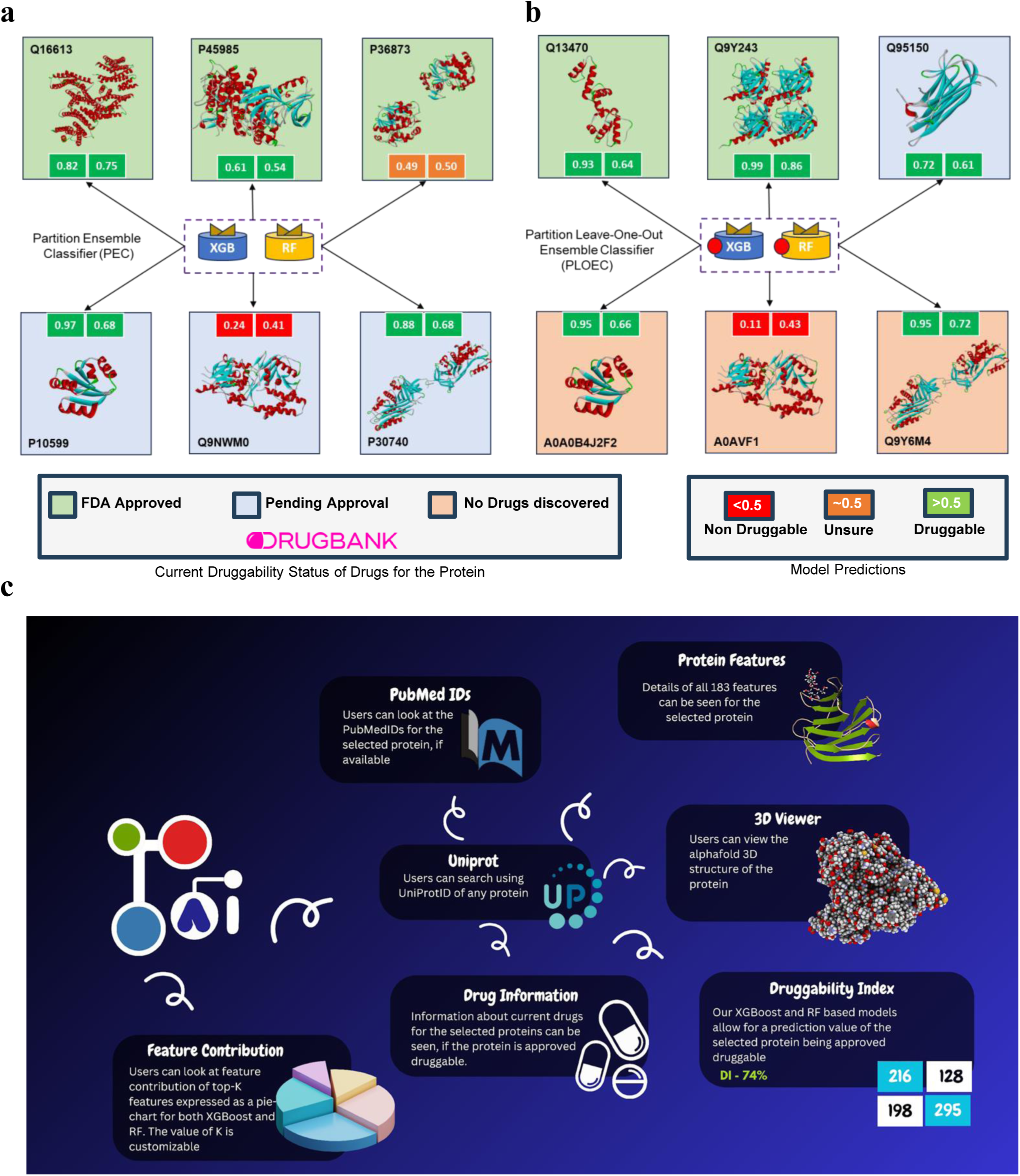
| Protein druggability status and predictions using PEC and PLOEC models and the layout of our tool DrugProtAI. The figure illustrates the druggability status of randomly proteins based on DrugBank data (dtd. 10 August 2024) and our Model predictions trained on druggability data (dtd. 16 March 2024) **a,** Six proteins initially under the investigational category. Top: three proteins now approved, with scores predicted by PEC using XGBoost (left) and Random Forest (right). Bottom: three proteins still pending approval, with two predicted as druggable by our model, indicating potential for future drug approval. **b,** Six proteins initially categorised as non-druggable. Top: two proteins now approved and one pending approval. Bottom: three remain non-druggable, but PLOEC identifies potential druggability for two. **c,** Depiction of the complete layout of our tool DrugProtAI.

### Overview of DrugProtAI

DrugProtAI is a robust web-based tool designed to assess protein druggability by integrating information from various biological knowledge bases and employing advanced machine learning techniques. By utilizing real-time data extraction from UniProt, DrugBank, PubMed, and AlphaFold, the application facilitates comprehensive protein druggability analysis, providing deeper insights into drug discovery and target validation for researchers. The primary objective of of DrugProtAI is to predict a protein’s druggability utilizing robust machine learning models, such as Partition Based Ensemble Models with Random Forest (RF), XGBoost (XGB), trained on 183 important biological characteristics. The top-ranking features are displayed in the form of interactive pie-charts allowing the users to visualize the feature contributions. The calculation of Druggability Index (DI) is one of the fundamental aspect of the tool that shall help the users (researchers, scientists, students, etc.) to prioritize the potential therapeutic targets. The tool also seeks to offer a rapid alternative solution for accessing all ongoing research pertaining to a specific drug target, thereby providing researchers with a unified platform for the preliminary stages of drug discovery. A schematic representation of DrugProtAI’s functionality is provided (**Fig. 3c**).

## Discussion

We have attempted to streamline the process of target identification by providing researchers with an advantage in interpreting the right drug candidates, saving significant time and effort in the drug discovery process. We present DrugProtAI, an easy-to-use web interface designed to facilitate the target selection process. DrugProtAI is built on the Partition Ensemble Classifier (PEC) and the Partition Leave-One-Out Ensemble Classifier (PLOEC), which enables robust, unbiased and accurate predictions of protein druggability, even with significantly skewed data. Data from DrugBank^18^ (https://go.drugbank.com/) revealed ∼84% of the proteins do not have any approved drugs against them whereas ∼16% have reported approved or investigational drugs against. Handling of such an imbalanced data has historically posed several complications, as also evident from the previous studies^7,9,12,19–22^ on protein druggability prediction. Our model employs a novel partitioning strategy, and we evaluate its effectiveness by measuring overall accuracy for multiple random seeds, as an equivalent substitute of cross-validation. The utility of the tool becomes apparent when tested on newly approved drugs, achieving around 80% accuracy in a blinded validation, thereby demonstrating its reliability. It outperforms current state-of-the-art tools. Furthermore, to the best of our knowledge, we are the first to identify the biophysical features that distinguish druggable proteins from non-druggable ones. We uphold a strong conviction that understanding these features will serve as a guiding force in selecting a target for drug identification when multiple protein targets associated with a particular disease are identified.

Kinases have long been reported as potential drug targets, and the FDA approval of Imatinib(DB00619) for cancer therapy has led to the emergence of a numerous kinase inhibitors in the field of oncology^23^. Our SHAP values and feature importance scores clearly highlight that proteins with kinase properties can be prioritized as potential therapeutic targets for designing inhibitors to treat and manage any disease. Additionally, hydrogen bonds play a crucial role in determining the stability of the proteins^24^ and shaping the structure of the proteins^25^. The secondary structure sheet is determined solely by hydrogen bonds^26^, and our results, along with the distribution pattern, show a distinct difference between distribution plots proteins with approved drugs and those without. The subsection of the UniProt^13^ providing information about modified residue (https://www.uniprot.org/help/mod_res) also appears to be among the top predictors, highlighting the importance of understanding protein modifications when selecting it as a drug target. Among the modified residues, glycosylation also emerges as a top classifier by the XGBoost partitioning model demonstrating strong concordance with the findings reported in literature^27–30^. It is quite obvious that drugs would tend to bind more effectively when protein structure tends to remain in a stable state, and our results reveal the instability index as one of the key regulators of the proteins to be druggable. Another important feature of proteins that has emerged as quite significant is aromaticity, as aromatic structures are known to play a role in forming protein-drug complexes and determining the binding affinity and specificity of drugs^31^. Among other important features, subcellular localizations such as the cell membrane and mitochondria are noteworthy. The proteins located on the cell membrane are particularly amenable to targeting for drug identification, which has led to the emergence of a new field focused on the cell surfaceome^32–35^. All these factors highlight the key features responsible for determining the druggability of proteins, which may be overlooked if only the sequence-specific properties of amino acids are considered, as is done in other contemporary studies. Thus, we posit that it may function as a valuable guide for investigators in the selection of specific drug targets.

The prediction becomes questionable when a particular data point is present in the training set itself. Therefore, we have also developed the partition leave-one-out method(PLOEC) to exclude that specific individual model which includes this protein in its training set from contributing to calculation of druggability index, thus ensuring no bias. Its utility becomes increasingly apparent when recent approved drugs are evaluated using this method, enabling us to achieve a level of accuracy surpassing that of existing tools. For instance, the recently approved drug targeting the protein Serotonin N-Acetyltransferase, a regulator of circadian rhythm^36^, has been assessed as druggable with exceptional ease using our method. Another protein within the mitogen-activated protein kinase (MAPK) family, integral to cellular proliferation and the promotion of metastasis in cancer cells^37^, has received approval for a drug. Our method, PEC, has effectively predicted its druggability in a favorable light. Similarly, the PLOEC method effectively identified tyrosine kinase non-receptor 1, an ubiquitin-binding protein associated with tumor growth^38^, as a viable druggable target, and this protein has recently been approved for therapeutic use. A similar result has been observed with another kinase, RAC-gamma serine/threonine-protein kinase. Our method indicates uncertainty regarding the serine/threonine-protein phosphatase PP1-gamma catalytic subunit, which may result from its features having less congruence with the currently recognized maximum druggable targets. Currently, for drugs that remain undiscovered and those awaiting approval, our tool effectively predicts outcomes, thus facilitating the selection of targets when multiple candidates are viable for addressing a specific disease. As a case in point, spermine oxidase, which is in the investigational phase, has been identified as non-druggable, whereas proteins such as SIK1B and casein kinase I isoform gamma-3, despite being classified as non-druggable, have been predicted to be druggable due to their potential associations with serine/threonine kinases. Testing on an independent protein set, such as Intraflagellar transport protein 56, similarly fails to secure a majority vote from all partitions as druggable. Consequently, our tool predicts a diminished likelihood of it serving as an effective therapeutic target. However, we would like to emphasize the importance of exercising caution when selecting a particular drug target based on the druggability index. For instance, while the protein complement factor D has recently received approval for a drug, our tool does not classify it as druggable. Now, the primary reason might be that complement factor D remains in its zymogen form and gets converted to active form by MASP-3^39^. Thus, it becomes evident that the properties may differ significantly from those of other proteins, potentially leading to incorrect interpretations by the tool. Delta-like protein 3, which is expressed in a range of diseases, including neuroendocrine prostate cancer, is currently being targeted by an antibody-drug conjugate^40^. However, our tool is unable to effectively analyze and predict its druggability. We firmly believe that incorporating data on the properties of proteins targeted by antibodies will further enhance the robustness of our tool. However, most of the recent proteins with approved drugs have been accurately predicted by the tool. Thus, we have thoroughly investigated the entire set of investigational proteins and provided a comprehensive guide for researchers to determine druggable targets

Most studies that have attempted to predict druggability do not provide a tool for experimental scientists to utilize, thereby limiting the scope primarily to theoretical scientists. Some existing tools only predict druggability while focusing exclusively on sequence properties. Thus, we developed an easy-to-use web interface that provides investigators with a comprehensive understanding of druggability, as well as the features that make proteins likely druggable in the near future. With a large number of differentially regulated proteins or genes at hand, a research scientist has the option to select a particular target for drug design, thereby increasing the likelihood of success in clinical applications. The recent pandemic has underscored the potential of drug repurposing^41^, and we are confident that it will substantially advance this strategy, ultimately conserving both time and financial resources. We are also providing the researchers full access to the publications pertaining to the particular protein used as a drug target.

DrugProtAI marks the inaugural step toward tackling the comprehensive spectrum of diverse druggability predictions, expanding the range to include other species, also including responses to antibodies, immunotherapies, antimicrobial agents, and drug resistance. With the growing availability of datasets and the advancement of novel algorithms and generative AI, we aspire to deliver more robust solutions to researchers in the near future.

## Methods

### Data Extraction and Pre-Processing

The dataset was developed by manually curating a dataset comprising of 20,434 human proteins and derived 183 features from the SwissProt-verified Uniprot database^13^. Proteins with sequence lengths exceeding 3,000 amino acids were excluded, thereby narrowing our dataset to 20,273 proteins. It was done to circumvent computational exhaustion of latent values. The feature sets were classified broadly into 10 distinct and well-defined categories. The features were derived from a diverse spectrum consisting of biological, physicochemical, and protein sequence-derived properties.

The dataset was thoroughly cross-referenced with DrugBank^42^. This integration facilitated the stratification of proteins into three distinct categories based on their associations with drug interventions. Proteins associated with one or more FDA-approved drug-targeted therapeutics are termed Druggable proteins. Proteins with one or more experimental or investigational drugs, currently undergoing clinical evaluations, are termed investigational proteins. Proteins with a lack of any known pharmacological agent, thereby not belonging to either of the aforementioned classes, are termed non-druggable proteins. An astounding 3.4% of proteins are associated with drugs that are still awaiting FDA approval. The dataset possesses an inherent skewness, with a large proportion of proteins remaining non-druggable, highlighting the challenges in drug discovery (**Fig. 1b**).

### Machine Learning Workflows and The Partition Method Employed to build a robust model to predict Druggability of Proteins

The machine learning (ML) techniques were employed to decipher the probable therapeutic action of a target protein using 183 extracted features. A novel metric has been devised, namely the ‘Druggability Index (DI)’ that can be used to determine the protein’s capacity to be druggable. DI can be calculated as the model’s output probability of predicting a protein to be druggable. 688 investigational proteins were excluded from our training dataset. The remaining dataset, comprising of 19,585 proteins, resembled a skewed distribution (2,636 druggable and 16,949 non-druggable). Model performance was evaluated on a held-out test set comprising 600 proteins from each of the druggable and non-druggable categories, ensuring robust evaluation. Consequently, the training dataset consisted of 18,535 proteins (2,036 druggable and 16,349 non-druggable). We provide the our entire dataset at **Supplementary Table 8**.

The substantial amount of imbalance in the dataset made it very crucial to handle it before implementing any kind of ML technique. The popular approaches like oversampling have been highlighted in different literature for handling imbalances^21^. Approaches such as synthetic oversampling like SMOTE^43^ and ADASYN^44^ are used to generate artificial samples of the minority class to balance the dataset. However, these methods lack transparency that the synthetic samples generated belong to true distribution of the minority class^45^. Another commonly used method undersampling may lead to loss of critical datapoints^45^.

We have used Random Forests^46^(Bagging or Bootstrap Aggregating) and XGBoost^47^ (Boosting), the two popular ensemble learning methods. It has been extensively used to handle imbalances in biological datasets and is increasingly being used to tackle this complicated issue of class imbalance in biological datasets^48,49^ thereby avoiding performance degradation. Mechanisms involved in bagging and boosting classifiers are depicted pictorially in (**Fig. 1c**). The primary reason for choosing these two ensemble methods is that bagging (used in Random Forest) trains several weak (less complex, small) models parallelly and independently, thereby reducing variance and improving model stability, whereas gradient boosting (used in XGBoost) sequentially trains models, where each model corrects the errors (or boosts the gradients) of the previous model, thereby minimizing bias and improving predictive accuracy. XGBoost has been implemented using the xgboost package and employs the Faster histogram optimized approximate greedy algorithm utilizing a default gbtree booster for the learning task of logistic regression for binary classification with a learning rate of 0.3, a max depth of 6, and a uniform sampling strategy with other hyperparameters at their default settings. Random Forest, on the other hand, is implemented using the sklearn package, uses the Gini criterion to measure split quality with 100 estimator decision trees, max depth is set to None, and remaining hyperparameters are at their default settings.

We have also utilised a genetic algorithm^50^ (GA) (**Supplementary Fig. 1**), a stochastic optimization methodology that boosts the selection of key features for druggability score prediction. Unlike, conventional deterministic methods like Recursive Feature Elimination (RFE), which are susceptible to being stuck in local minima, GA begins with a small diverse initial population of feature subsets. Three key genetic operations – crossover, mutation and selection is repeated until stopping criterion are met.

We then further proposed a **Partition Ensemble Classifier (PEC)**, which trains multiple ensemble models on different partitions of training data, while retaining the same feature set across partitions. The PEC classifier then performs an averaging over probabilities predicted by individual models followed by thresholding (>0.5) over the average probability to classify into druggable and non-druggable classes. Our hypothesis was that the ensemble of ensembles not only classifies well within the individual training sets, but also on the entire training data by collectively capturing subtle patterns and nuances. Each model might perform differently on other partitions, but the PEC’s final predictions were more balanced and are less prone to errors. The motivation behind our PEC design was backed up by our observation of poor performance on training ensemble models with unbalanced data. This partitioning ensured that each of the individual ensemble models was trained on a balanced dataset, while also ensuring that we are not ignoring critical properties of the majority class by using probability scores from each of the models for final prediction.

### Feature Importance and Interpretability in PEC model

The major limitation faced by contemporary tools is the lack of interpretable features depicting druggability. In any classifier algorithm, the features used to train the model are considered to be a true representation of the real-world scenario. However, this is not always the case, as noise in the data can also be inadvertently learned. Random Forest uses the mean decrease in Gini impurity to determine the important features^46^.On the other hand, XGBoost Classifier uses the Gain measure for feature scoring^47^. Our PEC model involved 6 models trained on 6 different non-druggable partitions. We reported the average feature importance scores for each model as the final feature score, ensuring a balanced consideration of their impacts. Though these built-in methods are well-suited for achieving higher performance metrics, they do not provide insight into how each feature affects the class predictions. Hence, we used SHAP (SHapley Additive exPlanations) values to gain better insights about impact on classes by different features^12,51^ . This technique is based on game-theoretic concepts, ensuring a fair distribution of credit (in this case, the prediction) among features^52^. The idea is that each feature’s contribution should be calculated by averaging over all possible permutations of feature inclusion, thereby ensuring fair credit by considering all possible feature interactions. SHAP values can be used to explain predictions for a given sample by summing the individual contributions from each feature. Average SHAP values across the entire training dataset provide a global picture of overall feature importance. For our PEC model, we reported the average of each model’s SHAP scores as the final Shapley values.

Given the model M and feature set 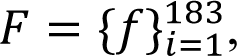, for a given input feature vector ξ ∈ ℝ^183^ corresponding to a protein, the final prediction is the sum of shapley values contributed by each of the feature 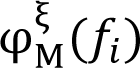, for feature *f_i_* plus a base value *B_M_* which is common across all inputs. The equation for model M’s final probability prediction 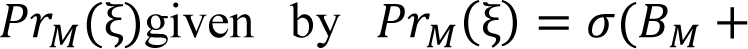 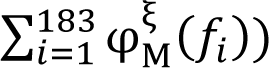. σ is a function that converts the shap values from the log-odds space to the probability space. Individual feature contribution is given by the following equation: 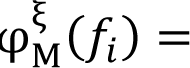 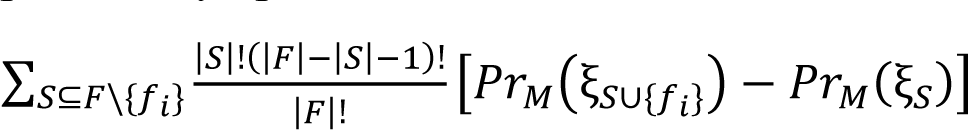 where ξ represents the feature values of ξ corresponding to the feature subset S. 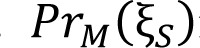 is calculated using a conditional expectation function by performing intervention on the S set of features given as: 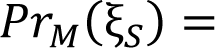 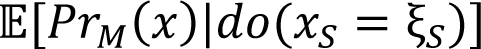. We used the TreeExplainer from python package shap, which was optimized for performance on tree-based models by computing shapley values in polynomial time^53^. For our proposed PEC model which was an ensemble of 6 models 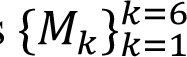, given an input feature vector ξ the shap value for a particular feature *f_i_* is given by average of shap value across 6 models: 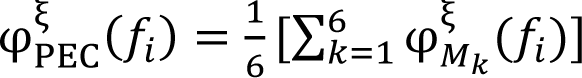.

### Quantifying Druggable Tendency by Druggability Index

Given a predictive model, the Druggability Index (DI) of a protein is defined as the probability of the given protein predicted by the model as druggable. The DI scores were separately obtained from the XGBoost and Random-Forest models following the aforementioned partition method to handle class imbalance. The PEC model in DrugProtAI is trained on the entire druggable and non-druggable protein sets. It completely excludes investigational proteins, to ensure that they are unseen during blind-validation.

We adopt separate strategies to calculate DI, depending on protein class.

1. For proteins belonging to the investigational and druggable class, the predictions were generated using the PEC (Partition ensemble classifier) model (Fig. 3a). The PEC model calculates DI value as the average of the prediction outputs of individual models trained on each partition of the dataset as proposed in the Partition method earlier. The prediction given by a PEC model *P*(*PEC*, ξ) in terms of prediction probabilities of individual models 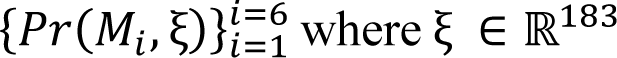 is the input feature vector corresponding to a non-approved druggable protein, is given by the equation: 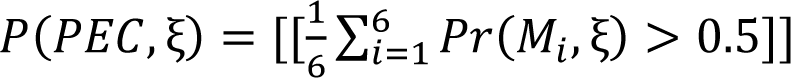 where [[*x*]] = 1 if x is true.
2. For proteins belonging to the non-druggable class, the predictions were generated using the PLOEC (Partition leave-one-out ensemble classifier) model (Fig. 3b). Every non-druggable protein is present in the training set in exactly one of the partitions. In the PLOEC model, we do not consider the partition containing this protein. The PLOEC model evaluates DI as the average of the prediction outputs of individual models trained on each of the remaining partitions. The prediction given by a PLOEC model *P*(*PLOEC*, ξ) in terms of prediction probabilities of individual models 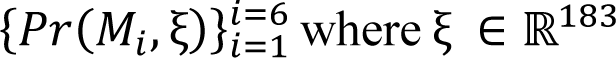 is the input feature vector corresponding to a non-druggable protein, is given by the equation: 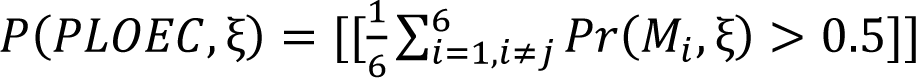 where *j* ∈ {1,2,3,4,5,6} be such that ξ ∈ *ND_j_* and [[*x*]] = 1 if x is true.

### Overview of our tool: DrugProtAI

DrugProtAI is powered by a Python-based Flask backend and features a dynamic application based on Javascript, while in-house styling was used for the layout of the webpage. Also, PostgreSQL was used for backend databasing and storing pre-loaded results. The interactive data visualization plots were created using the chart() JS function (https://www.jsdelivr.com/) . The 3D protein rendering tool leveraged 3Dmol.js (https://3dmol.csb.pitt.edu/).

The current version of DrugProtAI includes a dynamic website integrating the power of popular knowledgebases along with in-house curated ML models for comprehensive protein evaluation. We describe the functionalities and the working of the website:

**1. Search:** The landing page of the website allows the user to search for any protein based on its UniProt^TM^ ID^13^. This initiates multiple commands to retrieve pre-loaded and active information from multiple knowledgebases.
**2. Feature Contribution:** Our XGBoost (XGB) and Random Forest (RF) based models have been trained on 183 features and each of these features have certain contribution in the final decision. This contribution is displayed as a pie-chart available for both XGB and RF models. Users can view the contribution of top-K features (a total of K features having maximum contribution), where the value of K is user-driven. Note that since each of the two models have a set contribution for each feature, this is independent of the protein chosen.
**3. Druggability Index (DI):** The tool leverages the prediction results of both the XGB and the RF models to display the DI value of the protein selected. This feature is available for the proteins belonging to the non-druggable class or the non-approved druggable class. The proteins belonging to the class of approved-druggable proteins need not require the aforementioned metric.
**4. 3-D Viewer:** Users can view an interactive 3-D protein structure retrieved from AlphaFold^4^ of the protein selected.
**5. Protein Information:** The website retrieves information such as the functionality of the protein^13^. Along with this, each protein has a specified value of all the 183 features used for training. Users can view the values of all such features on the website. The PubMedIDs^54^ corresponding to the protein, if available – is displayed on the website.
**6. Drug Information:** The website displays the list of drugs attributed to the protein selected. This features is naturally available only for the proteins which have been approved druggable.

A schematic diagram showing the features of our tool can be seen (**Fig. 3c**)

## Code and Data Availability

All relevant code and data files for analysis can be found at https://github.com/Sachi-27/DrugProtAI.

## Author information

Corresponding authors (Lead contact)

Sanjeeva Srivastava*- Department of Biosciences and Bioengineering, Indian Institute of Technology Bombay, Powai, Mumbai 400 076; India

## Author Contributions

A.H., S.S.R. and S.S conceived and designed the project, A.H., S.S.R, S.B. performed data analysis, A.G. implemented the web tool, A.H., S.S.R, S.B., A.G. drafted the manuscript. S.S. edited the manuscript. All authors have approved the final version of the manuscript.

## Acknowledgements

We are grateful to the repositories and databases available which have been the major support to build the study. We are thankful to Mass Spectrometry Facility IIT Bombay (MASSFIITB). We also acknowledge Canva for providing scientific images. We would like to acknowledge Nirjhar Banerjee and Suhisna Dutta for helping us in designing the user interface of DrugProtAI. We would also like to thank Harshit Patil for helping us in drafting the manuscript. The study was supported through the MHRD-UAY Project (UCHHATAR AVISHKAR YOJANA) (UAY- (MHRD)), project #IITB_016 (2017) to S.S. MASSFIITB (Mass Spectrometry Facility at IIT Bombay) from the Department of Biotechnology (BT/PR13114/INF/22/206/2015). We also thank MERCK-COE (DO/2021-MLSP) for their extended support. AH was funded by the Ministry of Education, India through the Prime Minister’s Research Fellowship (PMRF) programme.

---

**Supplementary Fig. 1.**
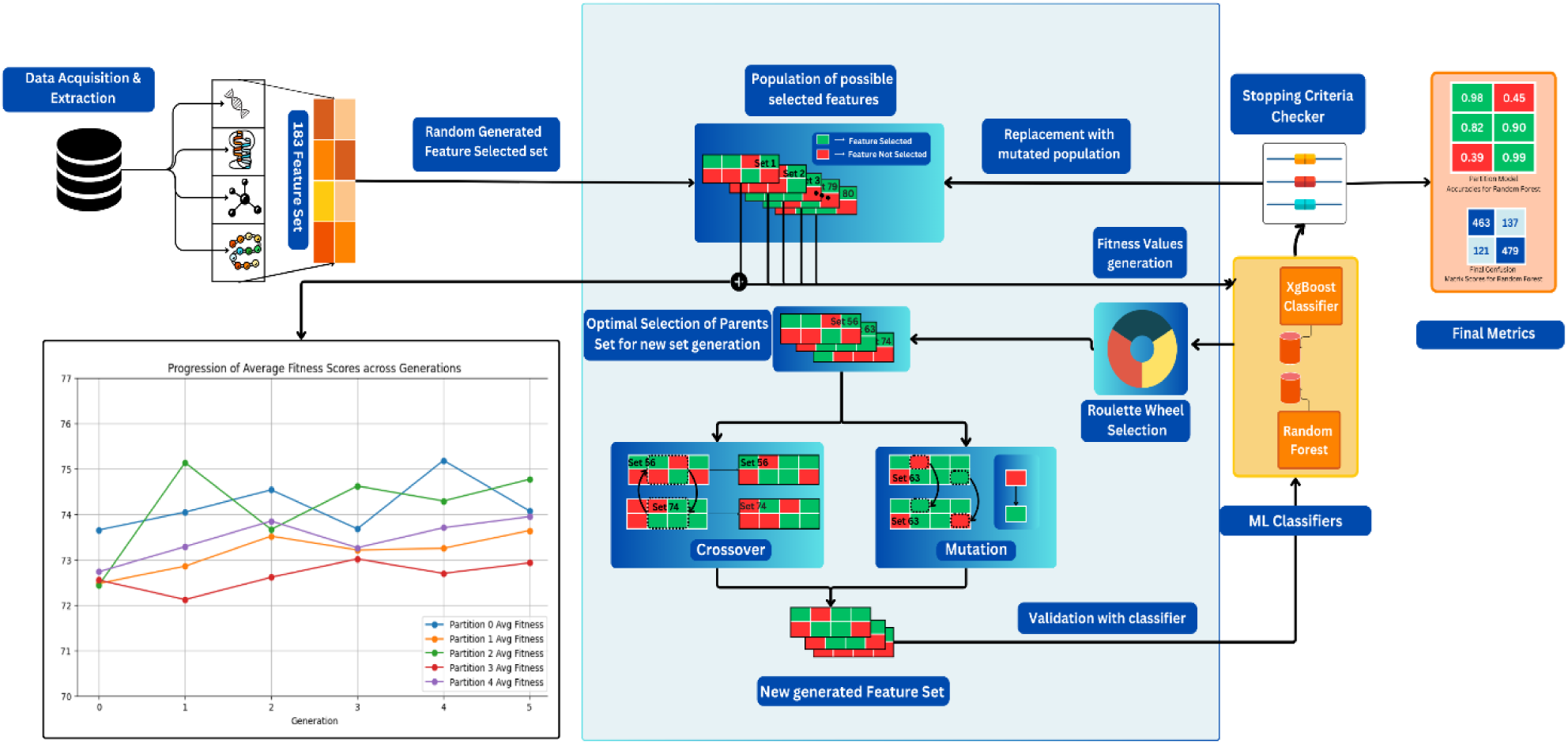
| Workflow of our Genetic Algorithm. The creation of a diversified initial population of feature subsets serves as the starting point for a GA-based feature selection procedure. Every subset offers a potential fix for the issue of feature selection. Three fundamental genetic processes are mostly used across a number of iterations: crossover, mutation, and selection. Initial operation of crossover involves combining features from two parent subsets to create new offspring subsets, facilitating the exchange of information between different feature sets. This process allows the algorithm to explore various combinations of features and inherit beneficial traits from parent solutions. Subsequently, mutation introduces variability into the feature subsets by randomly altering certain features. This randomness is crucial for preventing the GA from converging prematurely on local optima and helps in exploring a broader feature space. Finally, selection guided by a fitness function facilitated by ensemble model is designed to evaluate the predictive performance of each feature subset. To provide weightage to these fitness values, roulette wheel selection is used to choose feature subsets for reproduction based on their fitness scores. In this method, subsets with higher fitness scores are assigned a greater probability of being selected for the next generation, proportional to their performance.The GA process is continued until predefined stopping criteria were met, such as reaching a maximum number of iterations or achieving convergence i.e. in variance of the scores of the generated set . The plot **(left bottom)** shows plots of our fitness values across partitions for all 5 generation cycles.

**Supplementary Fig. 2.**
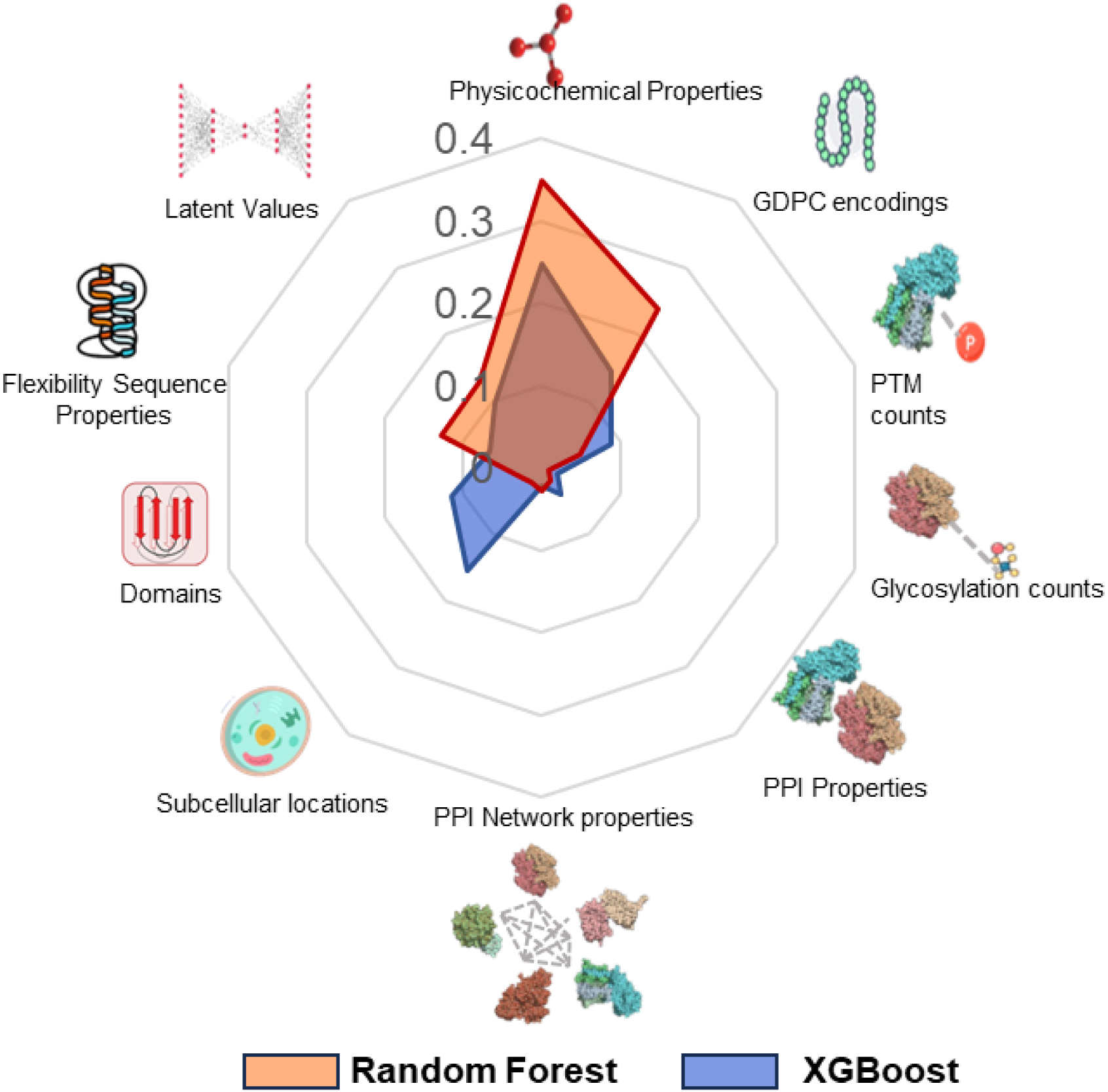
| Feature category wise importance scores for Partition Models. The figure illustrates the category wise total importance score calculated as sum of individual importance scores of every feature belonging to the feature category set for both XGBoost and Random Forest(RF) Partition Models, with feature scores calculated as partition average feature scores.

**Supplementary Fig. 3.**
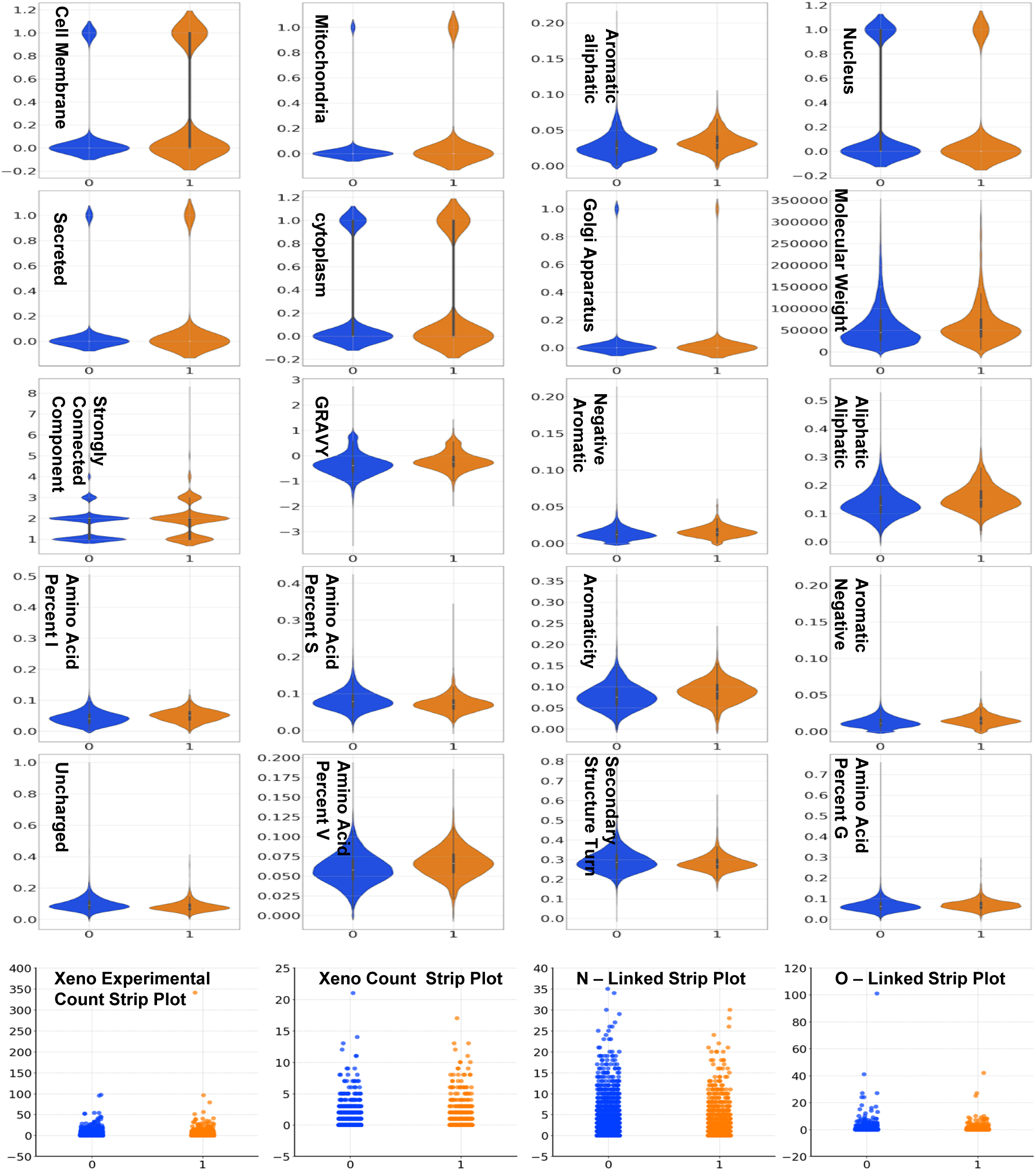
| Plots of top features in the intersection of Top 24 features of XGBoost and Random Forest based PEC models.

**Supplementary Table 3a.**
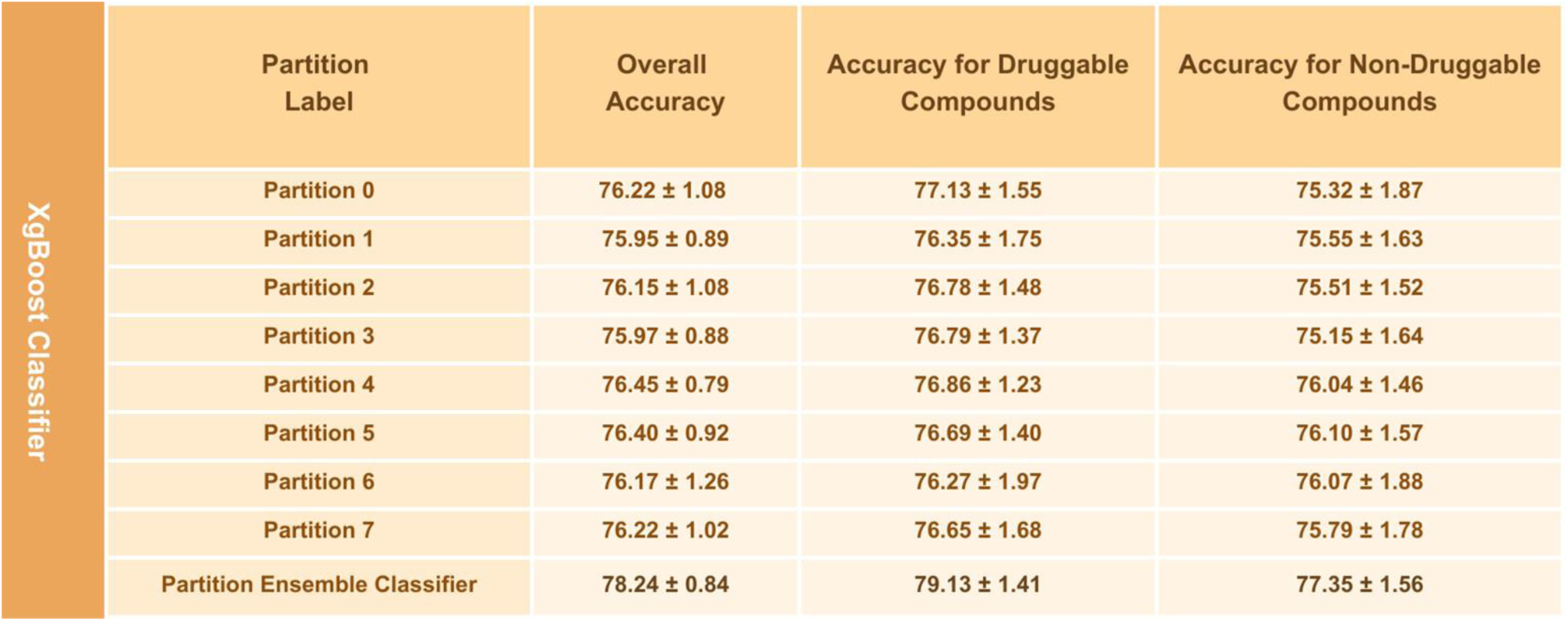
| Performance Metrics of the XGBoost Classifier using PEC Method Across Different Data Partitions

**Supplementary Table 3b.** | Performance Metrics of the Random Forest Classifier using PEC Method Across Different Data Partitions

| Random Forest Classifier | Partition Label | Overall Accuracy | Accuracy for Druggable Compounds | Accuracy for Non-Druggable Compounds |
| --- | --- | --- | --- | --- |
|  | Partition 0 | 74.85 ± 0.93 | 75.58 ± 1.58 | 75.58 ± 1.58 |
|  | Partition 1 | 75.32 ± 0.92 | 76.03 ± 2.09 | 76.03 ± 2.09 |
|  | Partition 2 | 74.86 ± 0.63 | 75.45 ± 1.61 | 75.45 ± 1.61 |
|  | Partition 3 | 75.16 ± 0.98 | 75.99 ± 2.08 | 75.99 ± 2.08 |
|  | Partition 4 | 75.29 ± 1.16 | 75.94 ± 1.69 | 75.94 ± 1.69 |
|  | Partition 5 | 75.28 ± 1.15 | 76.08 ± 1.82 | 76.08 ± 1.82 |
|  | Partition 6 | 75.36 ± 1.06 | 76.02 ± 1.79 | 76.02 ± 1.79 |
|  | Partition 7 | 75.27 ± 1.04 | 75.89 ± 1.16 | 75.89 ± 1.16 |
|  | PEC Prediction | 76.01 ± 0.93 | 77.54 ± 1.65 | 77.54 ± 1.65 |

**Supplementary Table 3c.**
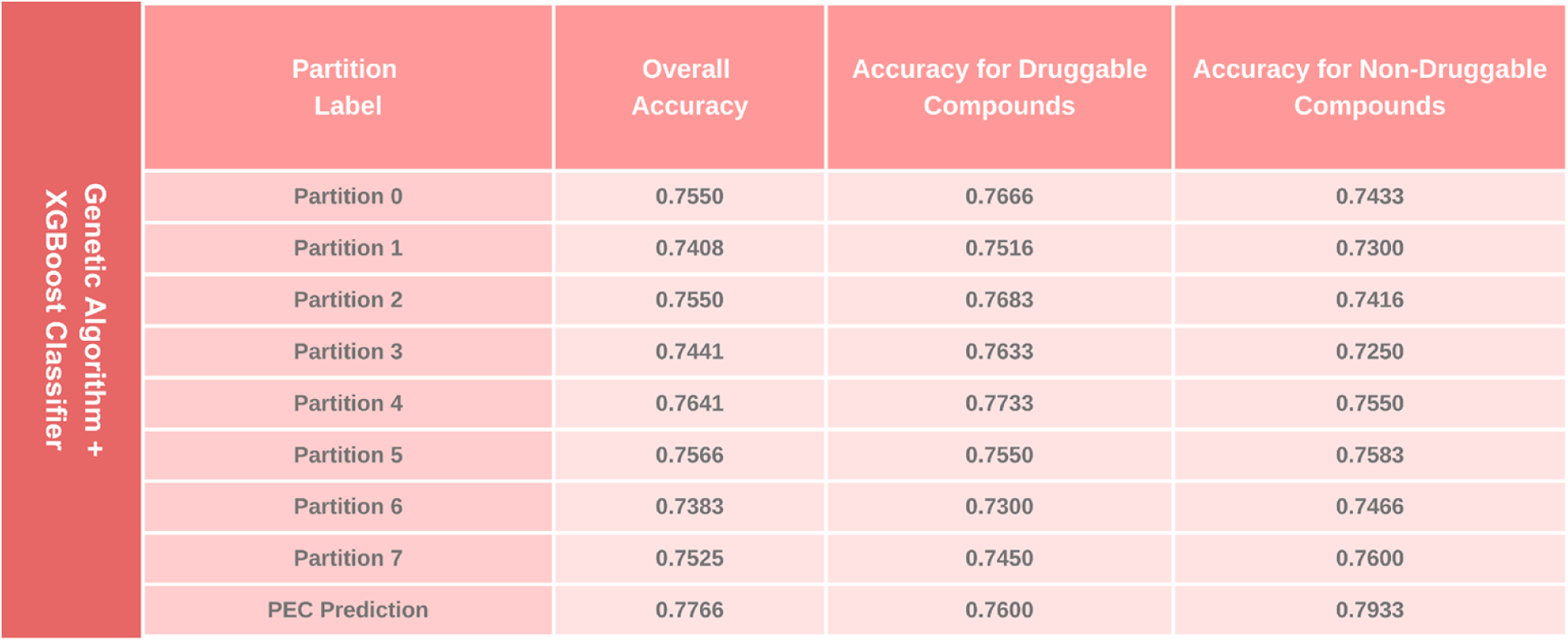
| Performance Metrics of the Genetic Algorithm Feature Selection with XGBoost Classifier using PEC Method Across Different Data Partitions

| Genetic Algorithm + XGBoost Classifier | Partition Label | Overall Accuracy | Accuracy for Druggable Compounds | Accuracy for Non-Druggable Compounds |
| --- | --- | --- | --- | --- |
|  | Partition 0 | 0.7550 | 0.7666 | 0.7433 |
|  | Partition 1 | 0.7408 | 0.7516 | 0.7300 |
|  | Partition 2 | 0.7550 | 0.7683 | 0.7416 |
|  | Partition 3 | 0.7441 | 0.7633 | 0.7250 |
|  | Partition 4 | 0.7641 | 0.7733 | 0.7550 |
|  | Partition 5 | 0.7566 | 0.7550 | 0.7583 |
|  | Partition 6 | 0.7383 | 0.7300 | 0.7466 |
|  | Partition 7 | 0.7525 | 0.7450 | 0.7600 |
|  | PEC Prediction | 0.7766 | 0.7600 | 0.7933 |

## References

1. Sun, D., Gao, W., Hu, H. & Zhou, S. Why 90% of clinical drug development fails and how to improve it? Acta Pharmaceutica Sinica B 12, 3049–3062 (2022).

2. Hajduk, P. J., Huth, J. R. & Tse, C. Predicting protein druggability. Drug Discovery Today 10, 1675–1682 (2005).

3. Batool, M., Ahmad, B. & Choi, S. A Structure-Based Drug Discovery Paradigm. IJMS 20, 2783 (2019).

4. Jumper, J. et al. Highly accurate protein structure prediction with AlphaFold. Nature 596, 583–589 (2021).

5. Abramson, J. et al. Accurate structure prediction of biomolecular interactions with AlphaFold 3. Nature 630, 493–500 (2024).

6. Sakharkar, M. K., Sakharkar, K. R. & Pervaiz, S. Druggability of human disease genes. Int J Biochem Cell Biol 39, 1156–1164 (2007).

7. Raies, A. et al. DrugnomeAI is an ensemble machine-learning framework for predicting druggability of candidate drug targets. Commun Biol 5, 1291 (2022).

8. Bull, S. C. & Doig, A. J. Properties of protein drug target classes. PLoS One 10, e0117955 (2015).

9. Bakheet, T. M. & Doig, A. J. Properties and identification of human protein drug targets. Bioinformatics 25, 451–457 (2009).

10. Finan, C. et al. The druggable genome and support for target identification and validation in drug development. Sci. Transl. Med. 9, eaag1166 (2017).

11. Shoombuatong, W., Schaduangrat, N. & Nikom, J. Empirical comparison and analysis of machine learning-based approaches for druggable protein identification. EXCLI J 22, 915– 927 (2023).

12. Charoenkwan, P. et al. Computational prediction and interpretation of druggable proteins using a stacked ensemble-learning framework. iScience 25, 104883 (2022).

13. Apweiler, R. et al. UniProt: the Universal Protein knowledgebase. Nucleic Acids Res 32, D115–119 (2004).

14. Halder, A., Verma, A., Biswas, D. & Srivastava, S. Recent advances in mass-spectrometry based proteomics software, tools and databases. Drug Discovery Today: Technologies 39, 69–79 (2021).

15. Vihinen, M., Torkkila, E. & Riikonen, P. Accuracy of protein flexibility predictions. Proteins 19, 141–149 (1994).

16. Cock, P. J. A. et al. Biopython: freely available Python tools for computational molecular biology and bioinformatics. Bioinformatics 25, 1422–1423 (2009).

17. Sikander, R., Ghulam, A. & Ali, F. XGB-DrugPred: computational prediction of druggable proteins using eXtreme gradient boosting and optimized features set. Sci Rep 12, 5505 (2022).

18. Wishart, D. S. et al. DrugBank: a comprehensive resource for in silico drug discovery and exploration. Nucleic Acids Res 34, D668–672 (2006).

19. Gong, Y., Liao, B., Wang, P. & Zou, Q. DrugHybrid_BS: Using Hybrid Feature Combined With Bagging-SVM to Predict Potentially Druggable Proteins. Front. Pharmacol. 12, 771808 (2021).

20. Arif, M., Fang, G., Ghulam, A., Musleh, S. & Alam, T. DPI_CDF: druggable protein identifier using cascade deep forest. BMC Bioinformatics 25, 145 (2024).

21. Jamali, A. A. et al. DrugMiner: comparative analysis of machine learning algorithms for prediction of potential druggable proteins. Drug Discov Today 21, 718–724 (2016).

22. Iraji, M. S., Tanha, J. & Habibinejad, M. Druggable protein prediction using a multi-canal deep convolutional neural network based on autocovariance method. Comput Biol Med 151, 106276 (2022).

23. Attwood, M. M., Fabbro, D., Sokolov, A. V., Knapp, S. & Schiöth, H. B. Trends in kinase drug discovery: targets, indications and inhibitor design. Nat Rev Drug Discov 20, 839–861 (2021).

24. Pace, C. N. et al. Contribution of hydrogen bonds to protein stability. Protein Science 23, 652–661 (2014).

25. Oklejas, V., Zong, C., Papoian, G. A. & Wolynes, P. G. Protein structure prediction: Do hydrogen bonding and water-mediated interactions suffice? Methods 52, 84–90 (2010).

26. Koch, O., Bocola, M. & Klebe, G. Cooperative effects in hydrogen-bonding of protein secondary structure elements: A systematic analysis of crystal data using Secbase. Proteins 61, 310–317 (2005).

27. Solá, R. J. & Griebenow, K. Glycosylation of therapeutic proteins: an effective strategy to optimize efficacy. BioDrugs 24, 9–21 (2010).

28. Costa, A. F., Campos, D., Reis, C. A. & Gomes, C. Targeting Glycosylation: A New Road for Cancer Drug Discovery. Trends in Cancer 6, 757–766 (2020).

29. Diniz, F. et al. Glycans as Targets for Drug Delivery in Cancer. Cancers 14, 911 (2022).

30. Smith, B. A. H. & Bertozzi, C. R. The clinical impact of glycobiology: targeting selectins, Siglecs and mammalian glycans. Nat Rev Drug Discov 20, 217–243 (2021).

31. Lanzarotti, E., Defelipe, L. A., Marti, M. A. & Turjanski, A. G. Aromatic clusters in protein– protein and protein–drug complexes. J Cheminform 12, 30 (2020).

32. Wei, J. et al. Profiling the Surfaceome Identifies Therapeutic Targets for Cells with Hyperactive mTORC1 Signaling. Molecular & Cellular Proteomics 19, 294–307 (2020).

33. Geri, J. B. & Pao, W. Elucidating the Cell Surfaceome to Accelerate Cancer Drug Development. Cancer Discovery 14, 639–642 (2024).

34. Ferguson, I. D. et al. The surfaceome of multiple myeloma cells suggests potential immunotherapeutic strategies and protein markers of drug resistance. Nat Commun 13, 4121 (2022).

35. Mandal, K. et al. Structural surfaceomics reveals an AML-specific conformation of integrin β2 as a CAR T cellular therapy target. Nat Cancer 4, 1592–1609 (2023).

36. Wadas, B. et al. Degradation of Serotonin N-Acetyltransferase, a Circadian Regulator, by the N-end Rule Pathway. J Biol Chem 291, 17178–17196 (2016).

37. Cargnello, M. & Roux, P. P. Activation and Function of the MAPKs and Their Substrates, the MAPK-Activated Protein Kinases. Microbiol Mol Biol Rev 75, 50–83 (2011).

38. Chan, T.-Y. et al. TNK1 is a ubiquitin-binding and 14-3-3-regulated kinase that can be targeted to block tumor growth. Nat Commun 12, 5337 (2021).

39. Dani, R. et al. Quantification of the zymogenicity and the substrate-induced activity enhancement of complement factor D. Front. Immunol. 14, 1197023 (2023).

40. Puca, L. et al. Delta-like protein 3 expression and therapeutic targeting in neuroendocrine prostate cancer. Sci. Transl. Med. 11, eaav0891 (2019).

41. Rodrigues, L., Bento Cunha, R., Vassilevskaia, T., Viveiros, M. & Cunha, C. Drug Repurposing for COVID-19: A Review and a Novel Strategy to Identify New Targets and Potential Drug Candidates. Molecules 27, 2723 (2022).

42. Knox, C. et al. DrugBank 6.0: the DrugBank Knowledgebase for 2024. Nucleic Acids Research 52, D1265–D1275 (2024).

43. Sowjanya, A. M. & Mrudula, O. Effective treatment of imbalanced datasets in health care using modified SMOTE coupled with stacked deep learning algorithms. Appl Nanosci 13, 1829–1840 (2023).

44. Munshi, R. M. Novel ensemble learning approach with SVM-imputed ADASYN features for enhanced cervical cancer prediction. PLoS One 19, e0296107 (2024).

45. Alkhawaldeh, I. M., Albalkhi, I. & Naswhan, A. J. Challenges and limitations of synthetic minority oversampling techniques in machine learning. World J Methodol 13, 373–378 (2023).

46. Breiman, L. [No title found]. Machine Learning 45, 5–32 (2001).

47. Chen, T. & Guestrin, C. XGBoost: A Scalable Tree Boosting System. (2016) doi:10.48550/ARXIV.1603.02754.

48. Salunkhe, U. R. & Mali, S. N. Classifier Ensemble Design for Imbalanced Data Classification: A Hybrid Approach. Procedia Computer Science 85, 725–732 (2016).

49. Liu, L. et al. Solving the class imbalance problem using ensemble algorithm: application of screening for aortic dissection. BMC Med Inform Decis Mak 22, 82 (2022).

50. Katoch, S., Chauhan, S. S. & Kumar, V. A review on genetic algorithm: past, present, and future. Multimed Tools Appl 80, 8091–8126 (2021).

51. Lundberg, S. & Lee, S.-I. A Unified Approach to Interpreting Model Predictions. Preprint at 10.48550/ARXIV.1705.07874 (2017).

52. Sun, M. W. et al. Game theoretic centrality: a novel approach to prioritize disease candidate genes by combining biological networks with the Shapley value. BMC Bioinformatics 21, 356 (2020).

53. Lundberg, S. M. et al. From Local Explanations to Global Understanding with Explainable AI for Trees. Nat Mach Intell 2, 56–67 (2020).

54. White, J. PubMed 2.0. Med Ref Serv Q 39, 382–387 (2020).

